# Natural avirulent *Shigella boydii* strain in the Brazilian Amazon lacks major virulence genes and present Type II, Type III and Type VI Secretion Systems

**DOI:** 10.1101/459701

**Authors:** Paula Taquita Serra, João Victor Verçosa, Ruth Moura de Souza, Paloma Inessa de Souza Dantas, Alan de Oliveira Rezende, Ana Paula Miranda Barros, Aline Rubens de Souza, Marcelo Ribeiro Alves, Marcelo de Souza Fernandes Pereira, Antônio Balieiro, Tainá Raiol, Luiz André Moraes Mariúba, Milton Ozório de Moraes, Sabrina Epiphanio, Najla Benevides Matos, Adolfo José da Mota, Gemilson Soares Pontes, Paulo Franco Cordeiro de Magalhães Júnior, Marcus Vinícius Guimarães de Lacerda, Paulo Afonso Nogueira, Patrícia Puccinelli Orlandi

## Abstract

**Background:** Among *Shigella* species, *Shigella boydii* has always displayed a smaller role to the overall *Shigella* burden, frequently placed at third in epidemiological studies and described as restricted to Southeast Asia. Here we characterize an *S. boydii* isolated from an epidemiological study enrolling 1,339 Brazilian children from the Amazon region, in which *Shigella* species solely was the fourth cause of bacterial diarrhea. *S. boydii* strain 183 was isolated from rotavirus co-infected children with acute diarrhea. Here we aimed to characterize this strain regarding virulence and, immune response in a pulmonary model.

**Methods:** An *in vitro* HEp-2 epithelial cell invasion assay was used to compare the invasive phenotype of *S. boydii* strain 183 with clinical and highly virulent *S. flexneri* strain, both isolated from Brazilian children. A murine pulmonary model was performed to assess lung damage by histopathological analysis. mRNA expression of immune response key genes was retrieved by multiplex real-time PCR and correlations were obtained by network analysis. Broad genome analysis was performed to confirm *S. boydii* identity and define its virulence profile.

**Results:** *S. boydii* strain 183 showed fewer invasion rates *in vitro* and tissue damage *in vivo* as compared to virulent *S. flexneri* 201. When compared to a survival challenge in mice, *S. boydii* had 100% survival against 10% of virulent *S. flexneri*. Overall, mRNA immune gene expression suggests a protective response against *S. boydii* strains 183, in contrast to the inflammatory response induced by the virulent *S. flexneri* strain 201. Network analysis with *S. boydii* strain 183 displayed IFN-γ protagonism, contrasting with the correlations centralized on TNF-α by the virulent *S. flexneri* strain 201. The genome showed a lack of effector proteins and enterotoxins in *S. boydii* strain 183, and sequencing analysis of *Ipa* invasins revealed mutations at functional sites. This avirulent *S. boydii* strain 183 presents the Type II Secretion System, T6SS, in addition to T3SS.

**Conclusions:** In addition to causing no disease, *S. boydii* strain 183 lacks effector proteins and enterotoxins. The presence of T6SS additional secretion system could provide an advantage to establish this strain among commensal bacteria.

**AUTHOR SUMMARY:** The *Shigella* genus is a human pathogen responsible to shigellosis and remains one of the significant causes of morbidity and mortality in children under five years old. This genus has four species, *Shigella flexneri*, *Shigella sonnei*, *Shigella boydii*, and *Shigella dysenteriae*. *S. flexneri* and *S. sonnei* are the most common in the worldwide infections; *S. dysenteriae* is rarely found, and *S. boydii* is responsible for 1% of the infections and is known to be restricted to Southeast Asia. Once *S. boydii* have a relatively small role in global *Shigella* disease, there are few studies regarding its virulence and mechanisms. Here we characterize an *S. boydii* isolated from Brazilian children from the Amazon region, and aimed to describe this strain regarding virulence. It is known that *Shigella* species use the Type 3 Secretion System (T3SS) to invade and colonize the human intestine. We found in *S. boydii* the presence of Type 2 Secretion System (T2SS), Type 6 Secretion System (T6SS), in addition to the T3SS. The T6SS have been described in *S. sonnei* only, granting a competitive advantage against *S. flexneri* mixed cultures. The presence of T6SS additional secretion system could provide a benefit to establish this strain among commensal bacteria.

## BACKGROUND

The *Shigella* genus is a Gram-negative, non-motile, facultative anaerobic human pathogen responsible for shigellosis and is a significant cause of morbidity and mortality in childhood and immunocompromised patients. Based on the structure of the lipopolysaccharide O-antigen, *Shigella* genus encompasses four subgroups: *Shigella flexneri*, *Shigella sonnei*, *Shigella dysenteriae*, and *Shigella boydii*, each composed of different serotypes (1), and their epidemiological profile. Recently, GEMS (Global Enteric Multicenter Study) classified *Shigella* as a top pathogen responsible for moderate-to-severe diarrhea in sub-Saharan Africa and South Asia in children under three years of age. During GEMS study, from the 1130 *Shigella* isolates collected in 36 months only 5.4% (61/1130) were identified as *S. boydii* (Livio *et al*. 2014). Although this number appears to be a proportionally small contribution to the overall cases compared to the other three *Shigella* species, *S. boydii* still is a significant component of the *Shigella* burden (2).

Considering the burden of disease caused by *Shigella*, there are relatively few *Shigella* genomic studies (2). Recent discussions about how the virulence factors of *Shigella* works to cause disease (3) still reinforces the need of whole-genome sequencing and genotyping of dominant species to identify the crucial virulence factors and contribute to *Shigella* vaccine development. Regarding virulence determinants to cause shigellosis, *S. boydii* is described as having the lower number of cases, representing 1-2% of all isolates and confined to the Indian subcontinent. The lower number of cases and the supposed lack of clinical relevance were pointed as the reasons why there is little known about *S. boydii* virulence determinants and its association with disease.

Although *S. boydii* are typically described as rarely found outside of South-East Asia (1), new serotypes are being discovery outside the Indian subcontinent (4). Here we characterize a *Shigella boydii* isolated from an epidemiological study enrolling 1339 Brazilian children from the Amazon region, in which *Shigella* species were the fourth cause of bacterial diarrhea, and the incidence of *S. boydii* was similar to that of *S. sonnei* (5). This wild-type *S. boydii* strain was lacking major virulence genes, enterotoxins and SPATE genes, and was unable to multiply in the cells *in vitro* and provoke injury in a murine pulmonary model, although it was isolated from diarrheic feces of rotavirus-infected children. Our findings show this non-virulent *S. boydii* str. 183 containing the presence of Type 2 and Type 6 Secretion Systems, in addition to T3SS, and eliciting a potentially protective immune response after challenge in mice. Once *Shigella* infection usually occurs person-to-person, we believe this additional secretion system could provide an advantage to establish this strain among commensal bacteria, withal once it has a non-invasive phenotype it may not cause the disease. This dichotomy could be the reason why *S. boydii* is poorly found in shigellosis cases. The presence of T6SS among *S. boydii* species and this suggested commensal nature could be used in future probiotic applications against shigellosis, and its genetic content information could increase the efforts towards *Shigella* vaccines development.

## MATERIAL AND METHODS

### Bacterial strains, cell culture, and growth conditions

*Shigella* species identification were performed by pentaplex PCR (6). Wild-type *Shigella* strain 183 were identified as *Shigella boydii* and was selected based on absence in *Shigella* enterotoxins genes *set-1A*, *set-1B, sen/ospD3* and *evt*, and other virulence genes such as *ipaH*, *invE* and *virF* (5). For comparison purposes, wild-type *Shigella* strain 201 identified as *Shigella flexneri* was also selected as a known virulent specimen (5). Bacterial strains were grown in Luria Bertani, Miller media (Himedia, Mumbai, India) at 37°C. HEp-2 cell (Merck KgaA, Darmstadt, Germany) culture was performed as described elsewhere (7). Briefly, HEp-2 cells were cultured in DMEM/F-12 medium (Invitrogen, Life Technologies, USA), supplemented with 10% fetal bovine serum (FBS), at 37°C under 5% CO2 atmosphere. *Shigella flexneri* 5a M90T was used as a universal invasion standard for intracellular multiplication and mouse challenge assays.

### PCR-based typing of virulence determinants

The virulence determinants were identified by PCR-based typing with some modifications from the original studies (5,8,9). The primers for PCR-based typing are described in S1.

### Intracellular multiplication on virulence assay

For the virulence assay, all bacterial strains were grown in 3mL of LB broth at 37°C overnight. The optical density was quantified at 600nm using a BioDrop DUO (Isogen Life Science, UK) spectrophotometer, and adjusted to 3×10^8^ CFU. HEp-2 cell culture was performed as described elsewhere (7). Briefly, a cell suspension was adjusted to 1×10^5^ cells/mL, and 2mL were distributed into a 24-well plate with a sterile round coverslip in the bottom. After a 24-hour incubation at 37°C with 5% CO_2_ atmosphere, the monolayer was washed with 0.85% sterile saline, and 100μL of bacterial suspension was placed and incubated for 1 and 3 hours. The supernatant was removed, and each well was washed four times with PBS 1X (Phosphate Buffered Saline). One milliliter of DMEM-F12 supplemented with SFB 10% (Gibco^®^, USA) with 100μg/mL gentamicin was added and incubated for an additional hour to eliminate non-invaded bacteria. The coverslips were removed, and the monolayer was preserved with Bouin Fixation Solution (Sigma-Aldrich, USA) and stained with Kit Panotico (Laborclin, Brazil) according to the manufacturer’s instructions. The stained coverslips were placed over a microscope slide and examined under immersion oil with 100X lens. All assays were performed in triplicate.

### Ethics, Anesthesia and Euthanasia

All efforts were made to prevent undue stress or pain to the mice. The mice were humanely euthanized once they show the following clinical signs: lethargy, hypothermia, and difficulty of breathing. The mice were euthanized with ketamine (300mg/kg) (Vetbrands, Brazil) and xylazine (22.5 mg/kg) (Syntec, Brazil), and consciousness was checked by testing the pedal re ex, heartbeats and breathing movements. All experiments were performed in accordance to the ethical guidelines for experiments with mice, and the protocols were approved by the National Council for Control of Animal Experimentation, The study was approved by the Animal Ethics Committee of *Instituto Nacional de Pesquisas da Amazônia* (INPA), N^°^. 018/2015, in regards to animal manipulation and experimentation. The guidelines for animal use and care were based on the standards established by The Brazilian College of Animal Experimentation (COBEA).

### Intranasal Murine Infection

Once adult mice do not exhibit disease symptoms upon oral administration of virulent *Shigella* spp. We chose the pulmonary infection route. Eight‐ to ten-week-old female Balb/C mice were anesthetized intramuscularly with 100μL of a solution containing 8-15 mg/kg of xylazine (Anasedan, VetBrands, Brazil) and 60-80 mg/kg of ketamine (Dopalen, VetBrands, Brazil). The 1×10^8^ CFU bacterial suspensions were adjusted to 40 μL in 0.85% sterile saline and administered through an intranasal route (10). After 24 and 48 h, mice were euthanatized, and lungs were removed to proceed to RNA extraction. All experiments were performed in three independent experiments, each with five mice.

### Lung collection and histopathological analysis

Mice were euthanatized, and lungs from infected and non-infected mice were treated similarly. Lungs were separated in two halves: one half was frozen for RNA extraction, and the other was fixed in 10% formalin for further processing. Paraffin-embedded non-consecutive lung sections were stained with hematoxylin-eosin (HE) and examined using light microscopy (Zeiss-PrimoStar, Zeiss). For histological and morphometric analysis, lung sections were blindly examined.

### RNA Extraction and cDNA synthesis

Whole-cell RNA was extracted from lung tissue using TRIzol^®^ (Invitrogen, Life Technologies, USA) according to the manufacturer’s instructions. RNA was treated with RNase-free DNase I to eliminate contaminating DNA using the RNeasy kit (QIAGEN, Hilden, Germany). Subsequently, RNA was quantified on Nanodrop ND-1000 (Thermo Scientific, Massachusetts, USA) spectrophotometer, and integrity was analyzed by agarose gel electrophoresis. The complementary DNA (cDNA) was synthesized from 100ng of total RNA using oligo (dT) primers and superscript III following the manufacturer’s instructions (Invitrogen, Life Technologies, USA).

### Multiplex real-time RT-PCR and gene expression analysis

Multiplex real-time RT-PCR assays were conducted with Fluidigm equipment using the Biomark platform (Fluidigm, California, USA) as described elsewhere (11). Twenty-three pair of primers from immune response genes or cytokines genes in mice were included, pre-amplified with a mix in a conventional thermocycler using TaqMan PreAmp Master Mix (Applied Biosystems, Life Technologies, USA) with 200nM aliquots of each primer and 1.25mL of each cDNA in a final reaction volume of 5 mL for 14 cycles. The cDNA from the GAPDH2 and B2M genes were employed simultaneously as expression normalizers. Pre-amplified cDNA was diluted 1:5 in DNA suspension buffer and loaded in the Fluidigm IFC Controller HX. Each probe was placed in the TaqMan Gene Expression Master Mix with Eva Green I stained (Applied Biosystems, Life Technologies, USA) and amplified in the Biomark Microfluidic System. Sample quality was analyzed using the company’s software, and raw data were exported. The fluorescence accumulation data from duplicate real-time RT-PCR reactions from one experiment were used in gene expression analysis to fit four-parameter sigmoid curves to represent each amplification curve using the qPCR library (12) for the R statistical package version 2.922 (13). The relationship between differentially expressed genes and sample profiles was investigated by Bayesian infinite mixture model cluster analysis (14) and represented by a 2D heatmap and dendrogram.

### DNA Sequencing

Genomic DNA was extracted (15) and DNA libraries were built using the Nextera XT DNA Sample Preparation Kit (Illumina, San Diego, CA, USA); sequencing was performed with HiSeq2500 (Illumina, San Diego, CA, USA).

### Bioinformatics analysis

A total of 1,989,616 paired-end reads were generated for strain 183, and 3,093,765 for strain 201 using the Illumina MiSeq platform (Illumina, San Diego, CA, USA). Reads were analyzed and trimmed using FastQC version 0.11.5 http://www.bioinformatics.babraham.ac.uk/projects/fastqc/ and FastX trimmer version 0.0.14 http://hannonlab.cshl.edu/fastx_toolkit. The reads were assembled using SPAdes version 3.9.0 (16), analyzed with QUAST (17), and annotated using RAST (18). Strain 183 assembly produced 237 contigs, with a total length of 5,034,743 bp, the largest contig of 178,316 bp, N 50 of 58,735 bp, and G⫹C content of 50.71%. Strain 201 assembly produced 342 contigs, with a total length of 4,431,146 bp, the largest contig of 96,160 bp, N 50 of 32,288 bp, and a G⫹C content of 50.50%. The virulence genes analyses were carried out using an integrated online *Shigella* database (http://www.mgc.ac.cn/ShiBASE/). The genomes and the remaining reads were converted to tabular files by using the Bioperl toolkit (http://bioperl.org/howtos/Beginners_HOWTO.html). The *Shi*BASE virulence gene sequences were used in the analysis and only those sequences which maximal segment pair (MSP) by BLASTN were selected. Multi-copy genes related to mobile DNA, such as IS elements and bacteriophage were excluded. The virulence protein commission information was retrieved from the KEGG Pathogen server (http://www.genome.jp/kegg/disease/pathogen.html). The same was performed with enzyme commission information and metabolic pathway maps; both were extracted from the KEGG ftp server.

A volcano plot was constructed (19) for visual display of mRNA expression analysis results. Briefly, the scheme was created with data obtained from the t-test and ANOVA, calculating the unstandardized signal (e.g., log-fold-change) against the noise-adjusted/standardized signal (e.g., t-statistic or ‐log(10) (p-value) from the t-test). The volcano plot is basic and interactive for understanding gene expression, regularized using criteria based on t-test and ANOVA. Two perpendicular lines referring to gene expression of the standard M90T strain, left, indicate lower expression than standard data, and right line, above expression, a 2-fold increase. The horizontal line discriminated whether dot plots differed statically, below the line indicates no significant difference and above the line (*p<0.05*).

### Statistical and Networking Analysis

To model interactions between the genes evaluated in this study, correlations between gene expression levels were assembled for each clinical strain, as described elsewhere (20). For networking analysis, Cytoscape software version 3.0.1 (Cytoscape Consortium San Diego, CA, USA) was used to draw biomolecule networks of correlations between two expressed genes, calculated using the Spearman’s rank correlation coefficient. When gene expression levels compared two bacterium results, we used non-parametric Kruskal–Wallis tests, and for three bacterium analyses we used ANOVA and plotted using GraphPad Prism software version 5.0 (San Diego, CA, USA).

## RESULTS

### Bacterial selection and characterization of virulence determinants

Bacterial strains used in this study were isolated from an epidemiological study enrolling 1,339 Brazilian children from the Amazon region (5). In order to confirm the results of 16S rRNA, we performed the pentaplex PCR typing method (6). As results, we found *S. boydii* as the second most frequent species with 30% of all *Shigella* isolates (data not shown), being this results disconnected from worldwide literature (1,3). The high frequency of *S. boydii* identified lead us to inquire about virulence characteristics, invasion behavior and genetic content of this *Shigella* species in the Amazon region, once there is no data about its presence in literature.

Strain 183 was isolated from rotavirus co-infected children with acute diarrhea and was negative for *Shigella* enterotoxins *shET1A*, *shET1B* and *shET2* (Figure 1A) (5). The whole-genome sequencing was performed, and the species was confirmed as *Shigella boydii,* re-named as *Shigella boydii* str. 183 (NCBI accession # CP024694). As *S. boydii* is defined as a not able to cause shigellosis species (3), we selected a different *Shigella* species to suit as positive shigellosis control. Strain 201 was identified as *S. flexneri* in pentaplex PCR (data not shown) and in whole-genome sequencing, renamed as *Shigella flexneri* str. 201 (NCBI accession # CP024981). This strain was isolated from monoinfected children with shigellosis and carried the majority of the virulence genes (Figure 1A) (5). As both of the *Shigella* strains were wild-type bacteria, we also compared our results with *Shigella flexneri* 5a (M90T), an invasion standard. All strains studied had *ipaBCD*, *ial* and *uidA* marker genes, necessary for the invasive phenotype. An important finding was that strain 183 was negative for the *virF* marker gene (Figure 1A).

**Figure 1:**
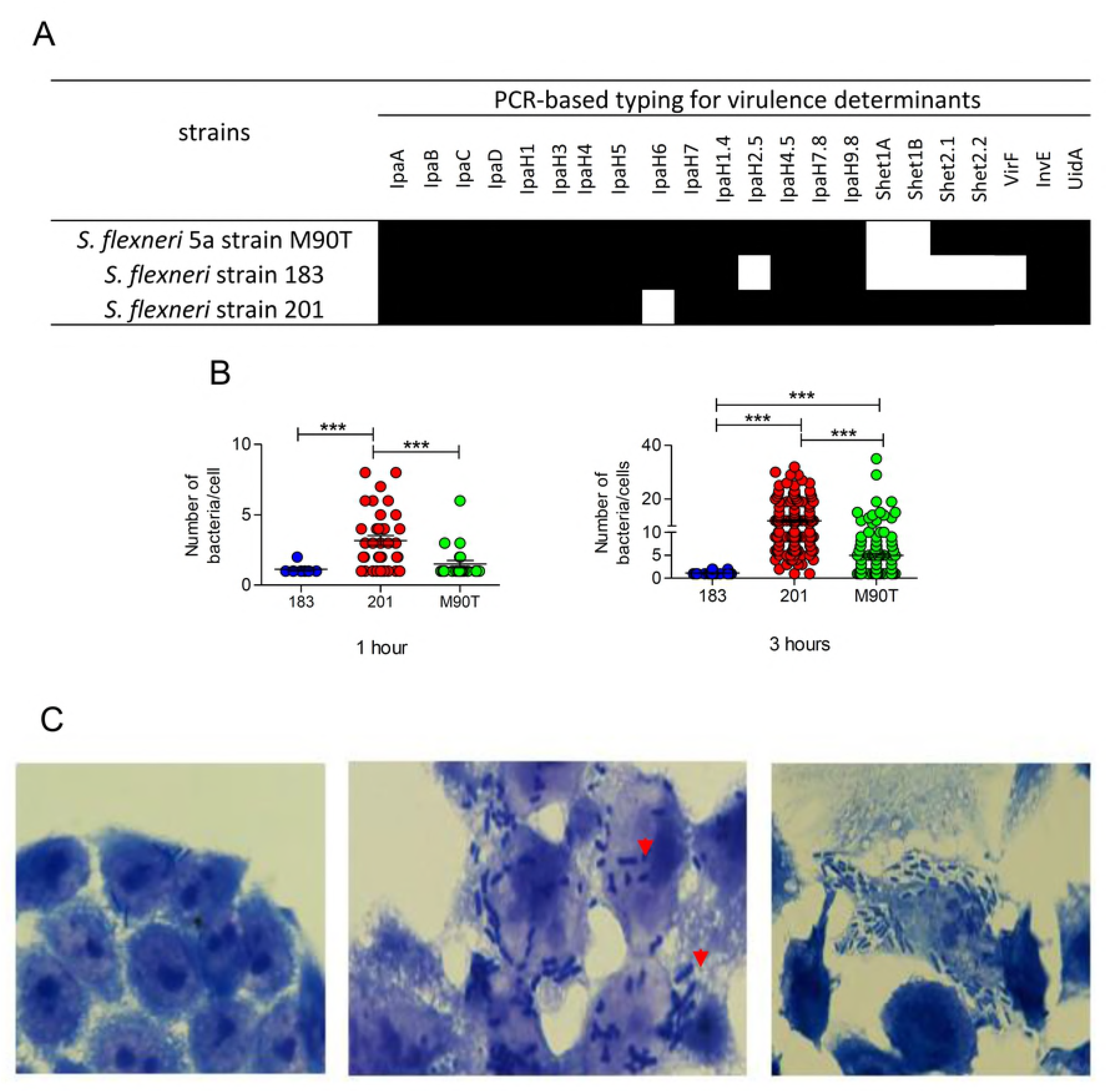
Virulence determinants of tested Strains: A) Classical PCR assay for detection of major pathogenicity determinants indicated strain 201 carried the majority of tested determinants. By our PCR assay, strain 183 did not contain *set1A*, *set1B* and *sen/ospD* genes. B) Comparison of intracellular multiplication ability among three tested strains was performed twice. The HEp-2 invasion assay was performed for 1 and 3 hours after cell contact. Graph shows the number of bacteria per cell. C) Photos show evolution of bacterial invasion and multiplication inside cells. Left panel: features of invasion; central panel: bacterial invasion and beginning of multiplication, bacteria in cell division (doublets); and right panel: bacterial invasion and intense multiplication in cytoplasm. Arrowhead: bacterial doublets in cell division.

We assessed the ability of the *S. boydii* str. 183 to invade epithelial cells (EC) and multiply in the cytoplasm *in vitro* using HEp-2 cell invasion assay (Figure 1B). We used the *S. flexneri* str. 201 as invasion reference, and while this bacterium displayed high invasiveness rates after one hour of cell contact, the *S. boydii* str. 183 had success in invading but not multiplying in the EC cytoplasm (Figure 1B). Surprisingly, this profile remained even after three hours of cell contact. The *S. boydii* str. 183 invasion rates did not increase, nor was their multiplication, suggesting that bacteria were unable to escape from the phagosome. Conversely, *S. flexneri* str. 201 had higher invasion rates, with several infected cells enclosing more than five bacteria per cell (*p<0,001*) after one hour of cell contact and substantially increasing after three hours with more than ten bacteria per cell (*p<0,001*).

To illustrate the *S. boydii* str. 183 behavior in the HEp-2 cell assay, we showed the evolution of *Shigella* invasion and multiplication inside the HEp-2 cell. After gentamicin treatment, lack of invasion features was assessed by the absence of bacterial cell inside the invasion vacuole (left panel; Figure 1C). The multiplication was noted by the occurrence of bacteria undergoing cell division, featuring bacterial doublets (central panel; Figure 1C). Intense bacterial amplification was observed in both virulent *S. flexneri* str. 201 and M90T (right panel; Figure 1C). *In vitro* HEp-2 cell invasion assay provided the invasion phenotype characterization of *S. flexneri* str. 201 as virulent invasive strain, while *S. boydii* str. 183 was less virulent or non-virulent, once it was not able to invade and multiply in the EC cytoplasm.

### Host Responses in mice infected with wild *Shigella* strains

Regarding invasiveness, we assessed the pathogenic potential of the *S. boydii* str. 183 in murine pulmonary infection. Eight‐ to ten-week-old female Balb/C were challenged by an intranasal route and survival rate was monitored for a hundred hours. Kaplan-Meyer analysis demonstrated that the survival rate of *S. flexneri* str. 201-infected mice were dramatically shortened compared to the *S. flexneri* 5a M90T standard (Figure 2A). In contrast, *S. boydii* str. 183-infected mice survived during all the monitored time (*p<0.0005*).

**Figure 2:**
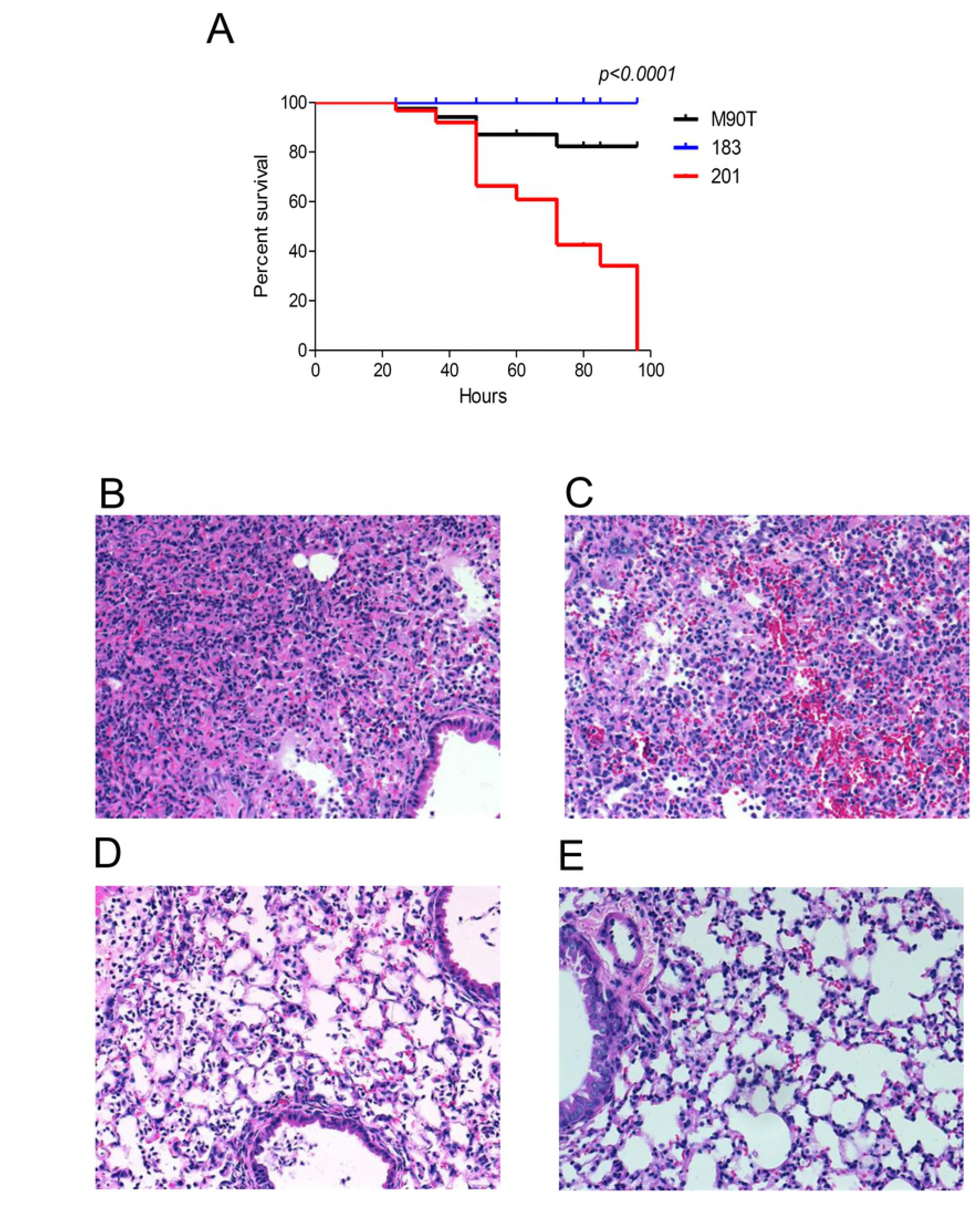
Immune response gene expression kinetics following challenge with wild *Shigella flexneri* strains. Eight‐ to ten-week-old female Balb/C were challenged intranasally with 1×10^8^ CFU of *Shigella* isolates; the first group (n=5) received the standard virulent strain *Shigella flexneri* strain M90T, the second group (n=5) was challenged with virulent strain 201, and the last group (n=5) received the avirulent strain 183. A) Results from two independent experiments each with five mouse specimens showed percentage survival measured using Kaplan-Meyer analysis and confirmed statistically that strain 201 was more virulent. B-E) Lung histopathological analysis from infected and non-infected mice. Lungs were fixed in 10% formalin for further processing with paraffin-embedded non-consecutive lung sections stained with hematoxylin-eosin (HE) and examined in light microscopy (Zeiss-PrimoStar, Zeiss). B) In *Shigella flexneri* strain 201-infected mice lung sections showed intense PMN infiltrates with the destruction of bronchial and alveolar epithelia, concomitant acute suppurative bronchiolitis, resembling the colitis that characterizes shigellosis. C) The same is observed in *Shigella flexneri* strain M90T-infected mice with the extended hemorrhagic area. D) In contrast, few infiltrates in lung occurred after infection with *S. boydii* strain 183 and any difference occurred in the alveolar epithelial morphology strain. E) Lung section from uninfected mice.

The pathology of *Shigella* infection in a pulmonary model shows bronchial and alveolar epithelia with concomitant acute suppurative bronchiolitis, resembling the colitis that characterizes shigellosis, as observed in both *S. flexneri* str. 201 and M90T (Figure 2B-C). Conversely, we found few infiltrates in the lung after infection with the *S. boydii* str. 183 and no changes in the alveolar epithelial morphology, as in uninfected mice (Figure 2D-E).

We assessed the immune response to define patterns of response-associated pathology and immunity. The gene expression of twenty-one immune-related genes were measured using multiplex real-time PCR mRNA gene expression. To assess the kinetics of gene expression, we gathered data from all strains in two-time points. Gene expression at 48 hours differed from that at 24 hours, and we found up-regulation of the Interleukin 1β (IL-1β), Interleukin 6 (IL-6), and NOD1 (Nucleotide-binding Oligomerization Domain 1), consistent with the premise of an innate immune response to bacteria (Table 1).

To determine the regulation range of the immune response genes, we measured qualitatively two-time points post-*Shigella* challenge, at 24 and 48 hours post-infection (Figure 3A). The majority of *S. boydii* str. 183-infected mouse genes were not expressed at 24h, a few were virtually down-regulated, and interestingly, genes TH1-like (Trihydrophobin 1) and MHC2-like (Major Histocompatibility Complex 2-like) were up-regulated in (Figure 3B). TH1-like is an integral subunit of the negative elongation factor complex (NELF), which inhibits signal transduction pathways associated with migration and adhesion cellular dynamics. MHC2-like is a general gene marker for the highly polymorphic H-2 complex. The up-regulation of TH1-like and MHC2-like genes in response to *S. boydii* str. 183 challenge could indicate a possible role of these genes in protective response, once infected mice had a 100% survival rate (Figure 3B).

**Figure 3.**
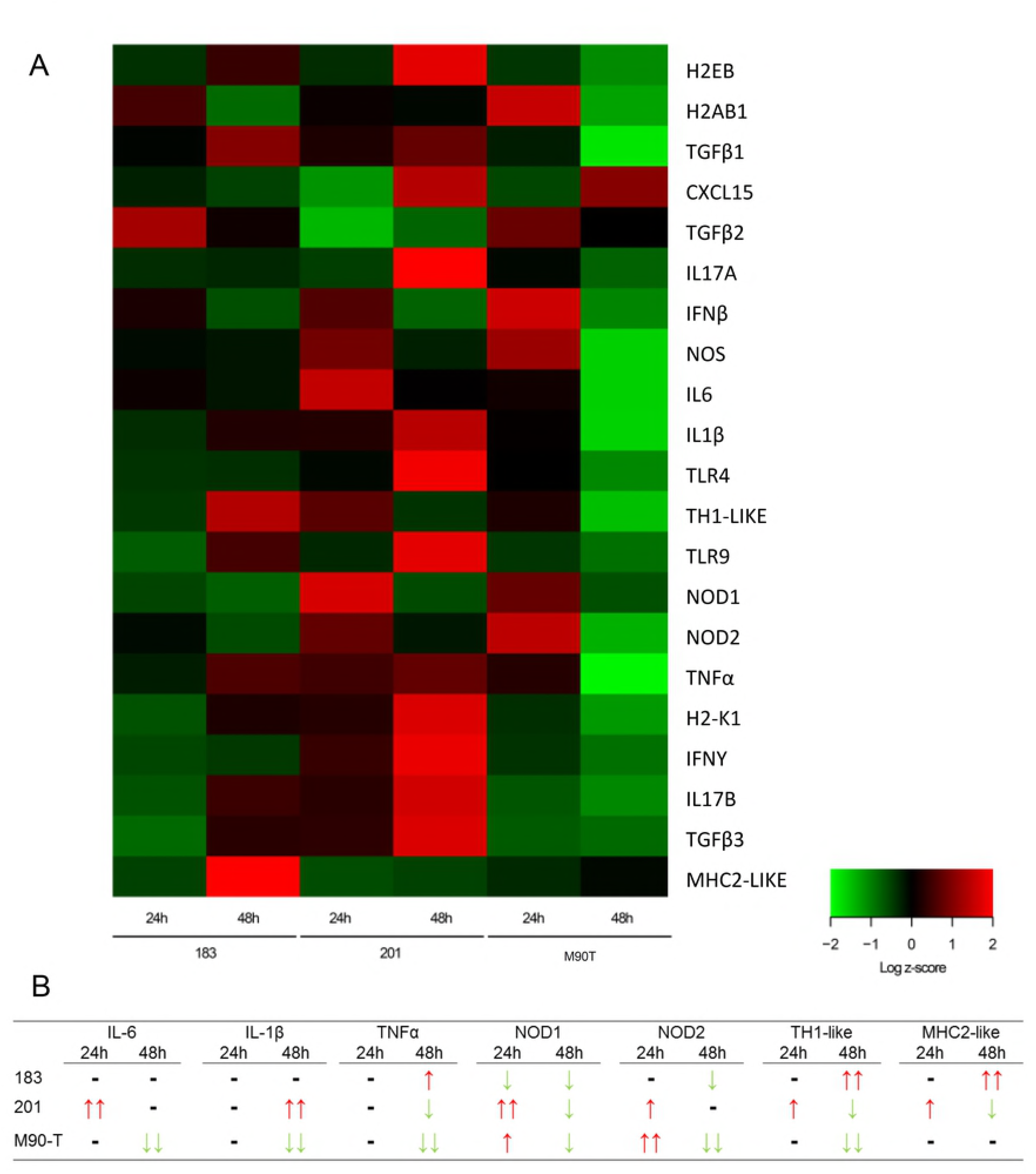
Immune response gene expression following challenge with wild-type *Shigella* sp. strains. Eight-to-ten-week-old female Balb/C mice were challenged intranasally with 1×10^8^ CFU of *Shigella flexneri* strain M90T (n=5), *Shigella flexneri* strain 201 (n=5) and *Shigella boydii* strain 183 (n=5). A) Multiplex real-time PCR was used for analyzing gene expression in the 24‐ and 48-hours post-infection groups. Gene expression of twenty-one immune response genes was determined in one experiment and grouped into two clusters (up‐ or down-regulation), as illustrated by a heat map. B) Considering only time points and not individual strain data, a summary of the kinetics of seven genes are compared for 24 – 48 hours (see Table 1). The color range illustrated by the heat map allows characterization of more or less regulated genes.

Immune response mRNA gene expression was compared across two days, and no gene was down-regulated below the normalizing GAPDH2 (Figure 4). *S. boydii* str. 183 induced up-regulation of several immune response genes at 24 hours in a way similar to virulent *S. flexneri* str. 201, and this profile was unchanged or reduced after 48 h (Figure 4). The expression of Tumor Necrosis Factor-alpha (TNF-α), Chemokine (C-X-C motif) ligand 15 (CXCL15) and IL-1β were induced slightly compared to the normalizer gene (Figure 4A). The innate, anti-inflammatory and cellular effector response genes of *S. boydii* str. 183 were up-regulated in 24h and reduced after 48h (Figure 4B-D). In *S. flexneri* str. 201 all pro-inflammatory cytokines were up-regulated in 48h, except IL-6 (Figure 4A). In meanwhile, gene expression of all other genes increased concomitantly with the innate response genes (Figure 4B). In contrast, all anti-inflammatory and cellular effector genes increased at 24h were reduced after 48 hours (Figure 4C-D).

**Figure 4.**
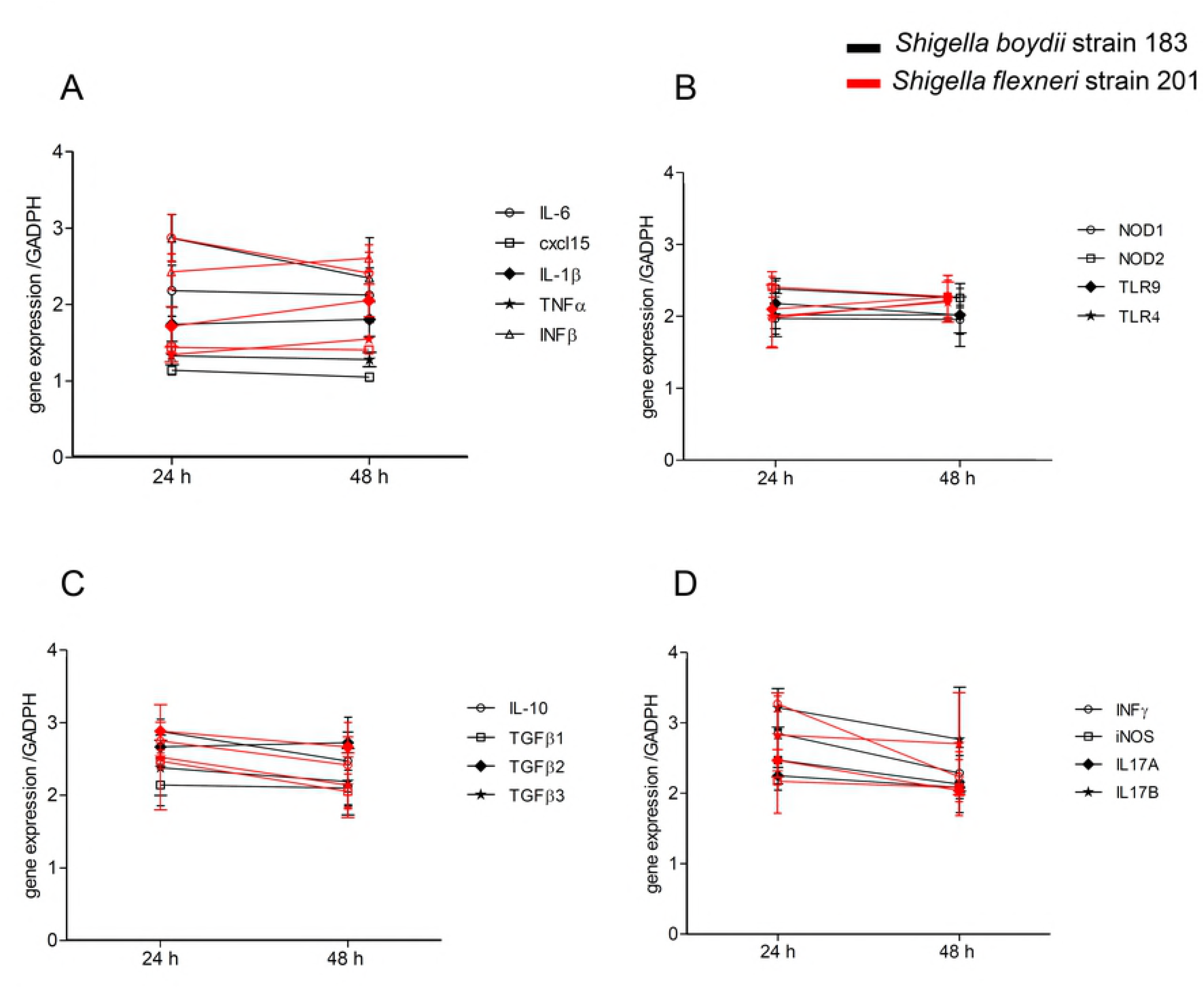
Immune response induced by *Shigella boydii* strain 183. Kinetics of immune response gene expression compared to those induced by *Shigella flexneri* strain 201 at two time-points: A) pro-inflammatory response genes; B) innate response genes; C) anti-inflammatory response genes; D) effector cellular immune response. Gene expression was quantified in groups of five and repeated once. Symbols and lines in red are *Shigella flexneri* strain 201. Symbols and lines in black are *Shigella boydii* strain 183.

Quantitative assessment of mRNA gene expression in 48h post-infection displayed *S. boydii* str. 183 inducing elevated expression of almost all immune response genes (Figure 5). Except Interleukin 10 (IL-10), Transformer Grown Factor Beta 3 (TGFβ3), inducible Nitric Oxide Synthase (iNOS) and Interleukin 17B (IL17B), *S. boydii* str. 183 had lower gene expression than virulent *S. flexneri* str. 201 and M90T standard.

**Figure 5.**
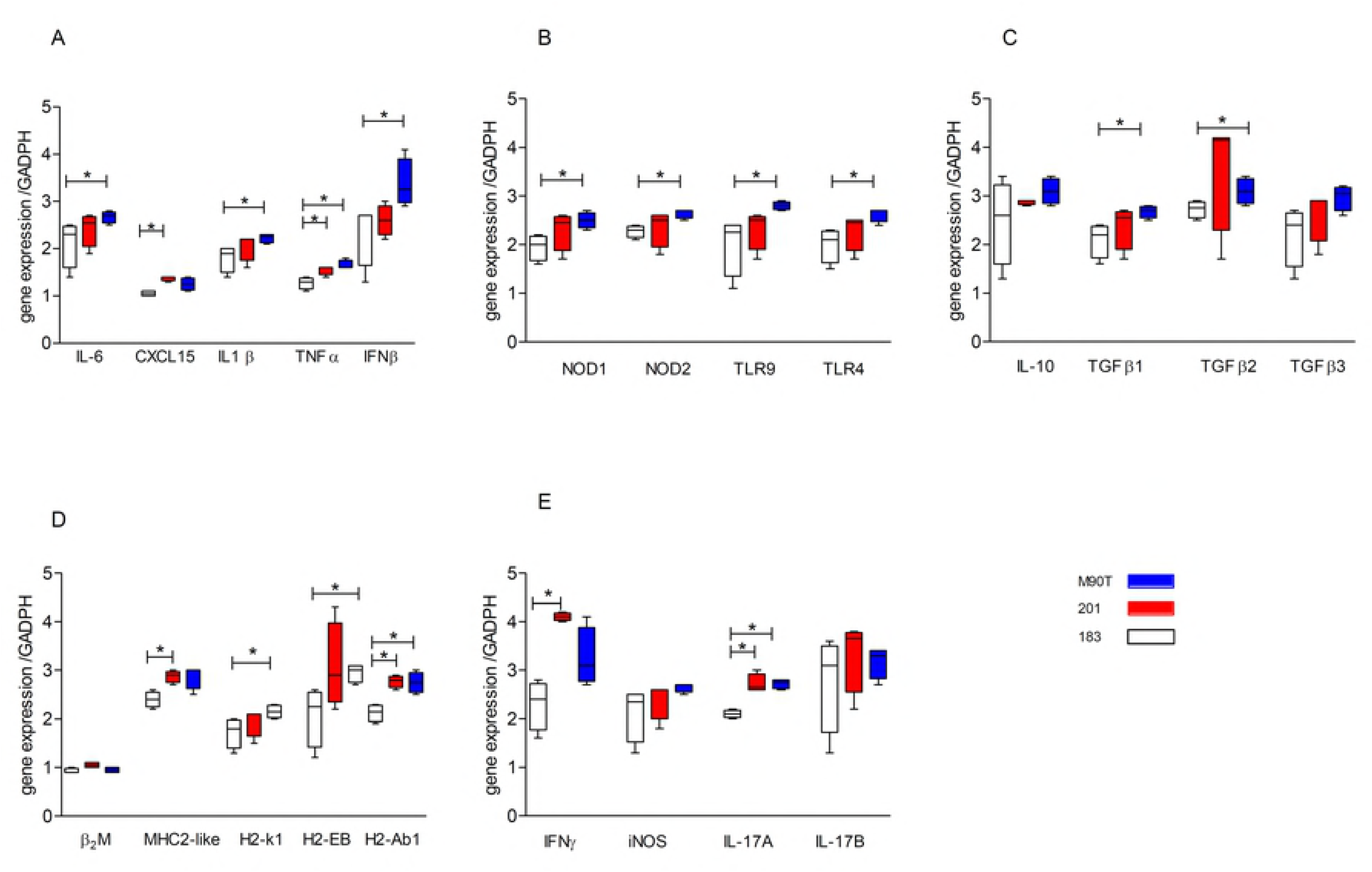
Quantitative assessment of gene expression after 48 hpi. Gene expression was compared among *Shigella boydii* strain 183, *S. flexneri* strain 201 and *S. flexneri* strain M90T after 48 hpi: A) pro-inflammatory response genes; B) innate response genes; C) anti-inflammatory response genes; D) effector cellular immune response. Gene expression was quantified in groups of five and repeated once. Comparison between two bacterial gene expression levels used non-parametric Kruskal–Wallis test. Asterisk indicated *p<0.05*.

In summary, although *S. boydii* strain 183-infected mice did not exhibit lung pathology, the strain induced a complete immune response without exacerbation, once adaptive response involves immune regulatory genes (Figure 5).

### Networking analysis is displaying two host response patterns

*S. boydii* str. 183 was considered a non-invasive isolate *in vitro*, and this phenotype was confirmed *in vivo*. On the other hand *S. flexneri* str. 201 had its invasive nature confirmed in both assays. A network analysis was performed with mRNA gene expression correlations of all genes tested at 48 hours post-challenge.

Interestingly, the host response network patterns changed with bacterial virulence (Figure 6). *S. boydii* str. 183 network revealed positive correlations of almost all genes with IFN-γ, while in *S. flexneri* str. 201 these correlations were towards TNF-α gene expression (Figure 6A-B).

**Figure 6.**
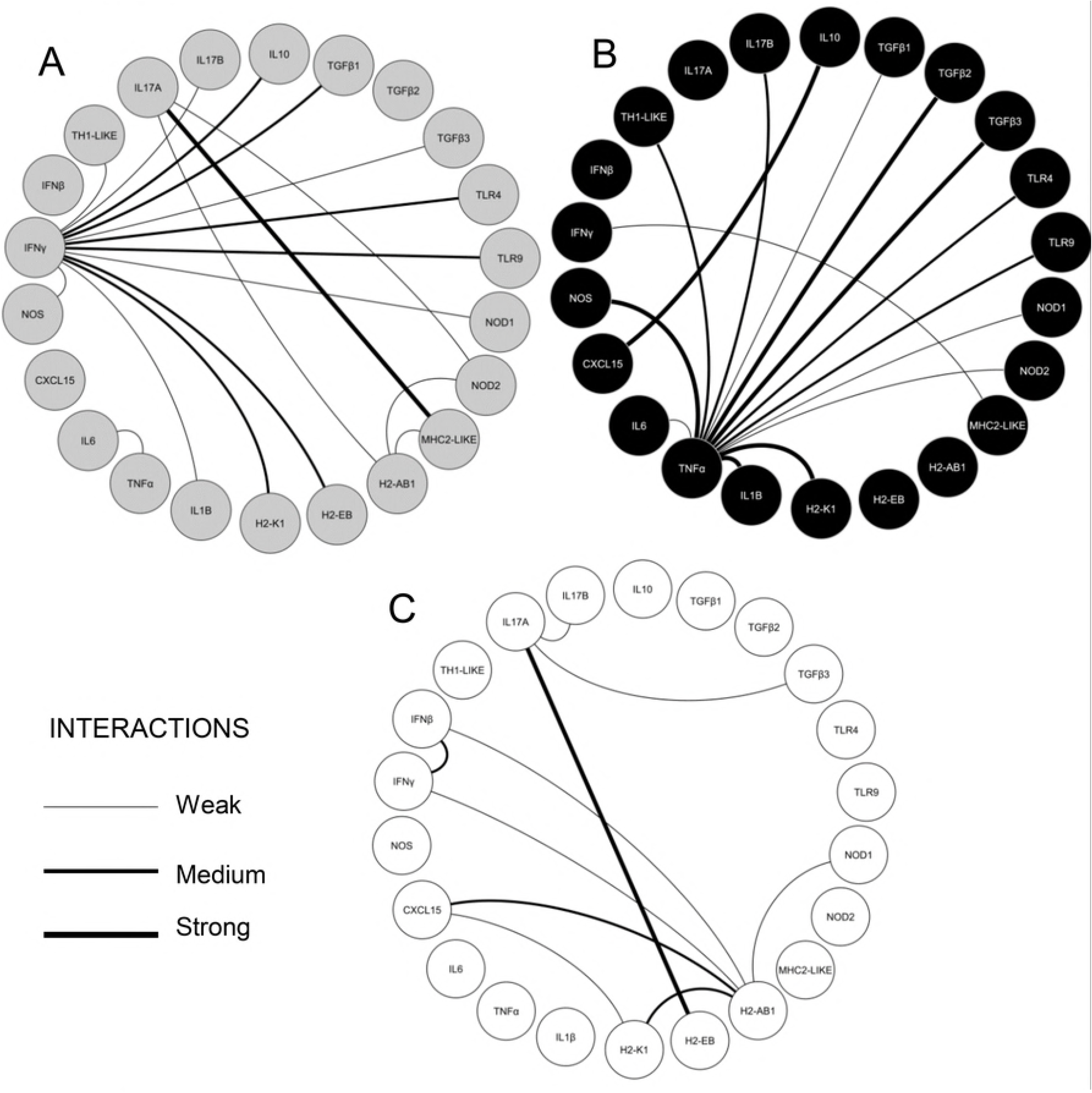
Networking of correlation between two expressed genes from host response patterns. Correlations between expressed genes were drawn to represent patterns of immune response at 48 hours post-infection. A) *Shigella boydii* strain 183; B) *Shigella flexneri* strain 201; C) *Shigella flexneri* strain M90T. The strength of interaction is represented as trace thickness. The networks of immune gene correlations were calculated using Spearman’s rank correlation coefficient. (thin trace: weak correlation, Spearman *r=0.7-0.79*; thinner trace = moderate correlation, Spearman *r=0.8-0.89*; thick trace = strong correlation, Spearman *r>0.9*). For network drawing, we used Cytoscape software version 3.0.1 (Cytoscape Consortium San Diego, CA, USA).

Conversely, we did not observe a clear gene expression pattern after M90T challenge (Figure 6C). This TNF-α converged network pattern for *S. flexneri* str. 201 is coherent with the lethal characteristic revealed by this isolate in a murine bacterial challenge with up-regulation of inflammatory mediators. Meanwhile, *S. boydii* str. 183 network pattern exposing IFN-γ protagonism is consistent with mice survival rate, suggesting a protective response.

### Genome comparative analysis and virulence content

To compare the genomic composition of both *S. boydii* str. 183 and *S. flexneri* str. 201 we performed a whole-genome sequencing. A map identifying a list of common gene systems was designed to access the virulence differences (Data Not Show). There are shared genes for secondary and energetic metabolisms and biosynthesis; gene systems associated with biochemical processes involving DNA, motility, transport, and cell signaling; and the type III secretion system (T3SS).

Conversely, *S. flexneri* str. 201 exclusively harbored gene systems associated with pili assembly, siderophores, and genes related to host interactions. In turn, *S. boydii* str. 183 harbored the Type 2 Secretion System (T2SS) and Type 6 Secretion System (T6SS) (Figure 7), and several gene systems associated with transport and membrane components.

**Figure 7:**
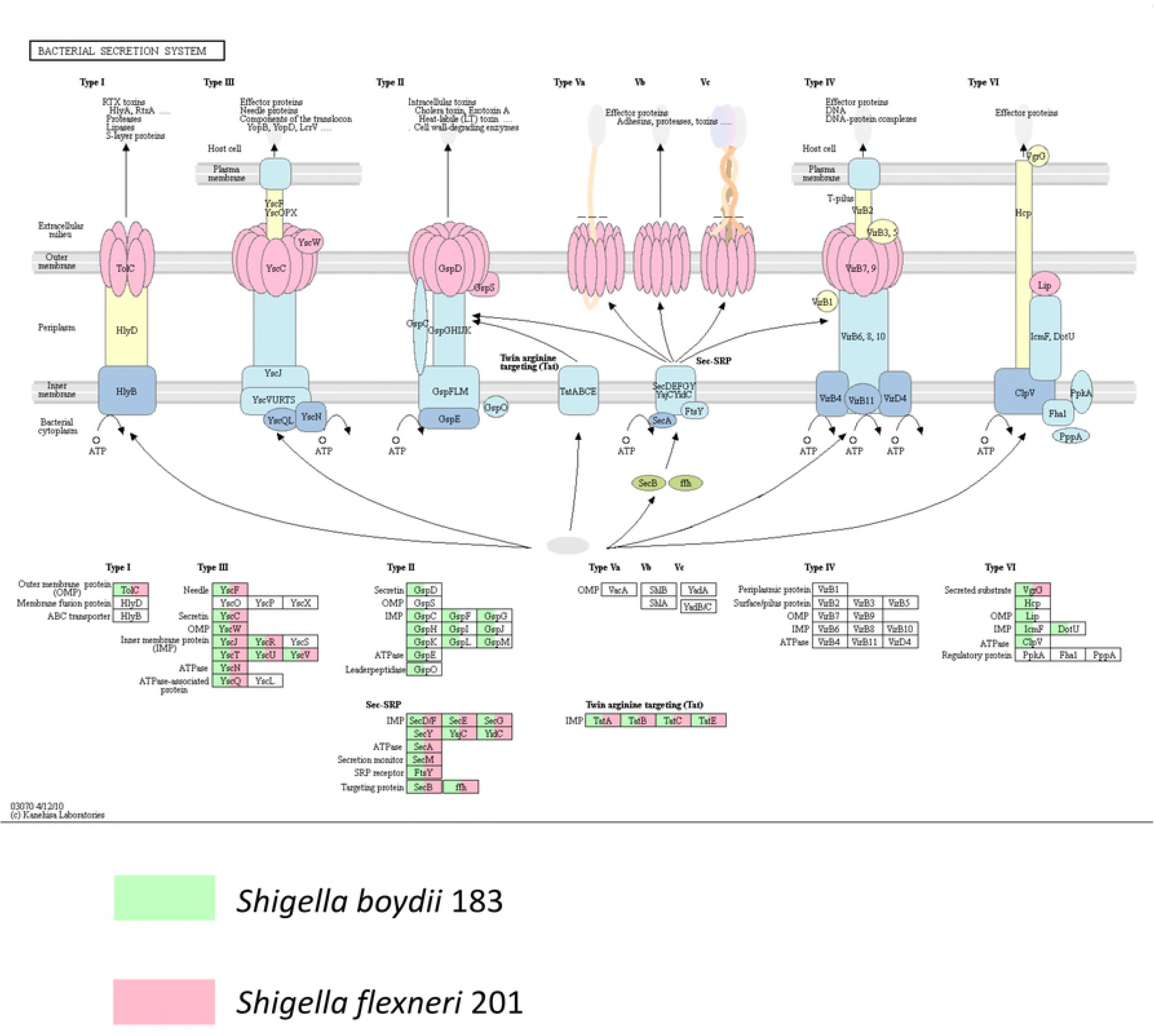
Secretion Systems of wild *Shigella* sp. *Shigella boydii* strain 183 genomes was compared with that virulent *Shigella flexneri* strain 201 at Secretion Systems possessing at KEGG. The color green indicates the gene presence to strain 183 and the pink color, to strain 201.

The virulome map showed a mosaic of virulence-related gene sets shared and exclusive of both bacteria (S2-5). In shared virulence-related genes, the virulome map displayed complete T3SS and a flagellar system apparatus. Curiously, *S. boydii* str. 183 harboring a set of genes for three alternative types of Gram-negative protein Secretion Systems that were absent in *S. flexneri* str. 201 (Figure 7 and S3). *S. boydii* str. 183 harbored an incomplete set of genes with similarity to components of the multi-protein complexes T2SS, the conjugation and translocation T4SS apparatus and the widespread specialized machinery T6SS, all involved in the delivery of toxins and effector proteins (Figure 8 and S3). Additionally, the virulent S. flexneri str. 201 harboring a set of effector proteins and enterotoxins encoded by *ospE1*, *ospE2*, *pic*, *sepA*, *set1A*, *set1B*, *vat*, *ospD3/senA*, which were absent in *S. boydii* str. 183 (S3).

**Figure 8.**
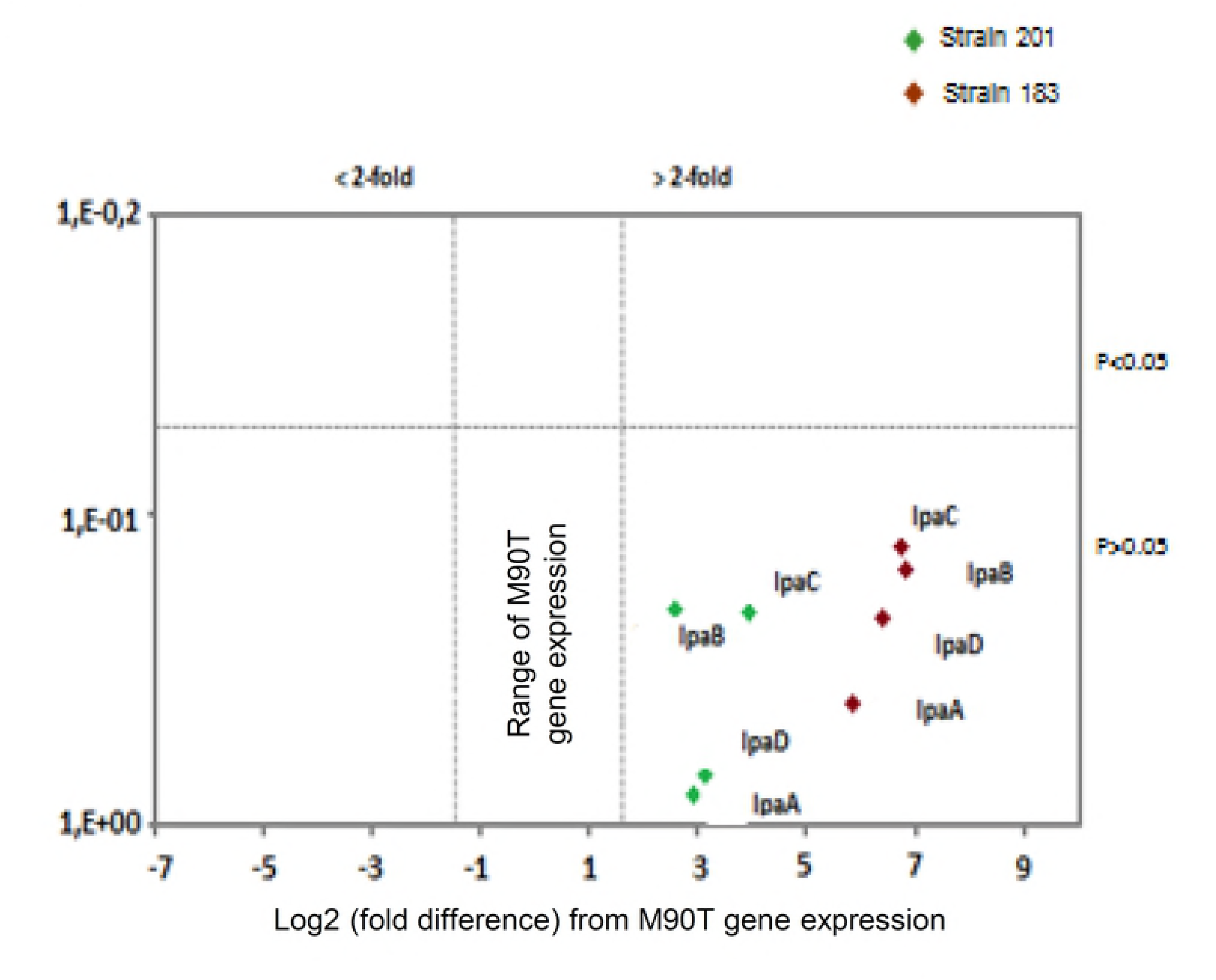
Balance of *ipaA, B, C* and *D* genes expression. Comparison of gene expression of major components of T3SS *Shigella*. After normalization, *ipaA*, *ipaB*, *ipaC* and *ipaD* expression levels in mice infected with strains 201 and 183 were compared to M90T strain-infected mice and represented as a volcano plot. The lower-right quadrant plotted genes with 2-fold higher expression than observed in M90T strain-infected mice (right dashed vertical borderline). The *ipaA*, *ipaB*, *ipaC* and *ipaD* levels did not differ and are plotted in same quadrant.

Both strains shared multiple genes with similarity to components of the Flg, Flh and Fli systems related to flagellum assembly (S4). Once again, *S. flexneri* str. 201 harbored a complete set of type 1 pili fimbrial systems, incomplete in *S. boydii* str. 183 (S5). Furthermore, both strains harbored incomplete gene sets of the regulator (Che and Vir systems), transporter (Ecp system), and antigen modification genes (Gtr and Csg systems). One relevant finding was the presence of the global transcriptional regulator gene (*virF*) in only the virulent *S. flexneri* str. 201. Ultimately, the virulence map showed several siderophore-mediated systems produced by pathogenic Gram-negative bacteria. Both strains shared Ent and Fep siderophore systems, similar to bacterial enterobactin and ferrisiderophore, respectively. *S. flexneri* str. 201 harbored Iuc-aerobactin and Sit system, and *S. boydii* str. 183 sheltered Irp and Ybt systems; all are iron-uptake microbial systems primarily found in Gram-negative bacteria (S5).

### Expression of T3SS genes does not explain distinct virulence behaviors

Although the virulome suggested that *S. flexneri* str. 201 virulence phenotype could be associated with its genetic content, the functional gene expression *in vivo* of this phenotype was still in question. Hence, we assessed the mRNA gene expression of components of T3SS, the primary system used by *Shigella* spp. to assembly invasion apparatus.

After normalization of *ipaA*, *ipaB*, *ipaC* and *ipaD* expression with housekeeping gene *rrsA*, gene expression levels of infected mice with *S. flexneri* str. 201 and *S. boydii* str.183 were compared to M90T standard and represented in a volcano plot (Figure 8). The assessment shows gene expression plotted in the lower-right quadrant. Although *ipaA*, *ipaB*, *ipaC*, and *ipaD* levels were expressed 2-fold highly than in M90T-infected mice (dashed vertical borderline), mice infected with *S. flexneri* str. 201 did not differ compared to those of *S. boydii* str. 183, constituting a prominent finding for the *Shigella*-host interaction.

### Mismatches in T3SS apparatus needle tip as virulence loss

The similar gene expression of *ipaA*, *ipaB*, *ipaC*, and *ipaD* by both bacteria suggests that non-invasive phenotype displayed by *S. boydii* str. 183 was not related to the absence of T3SS gene expression, but maybe to a possible polymorphism causing remodeling of the components structures. The *ipaA*, *ipaB*, *ipaC*, and *ipaD* amino acid sequence were searched for mismatches and 3D-structure *in silico* were predicted to determine whether a possible remodeling in these structures could be a factor associated with loss of virulence.

The *ipaA, ipaB* and *ipaC* genes of *S. boydii* str. 183 and *S. flexneri* str. 201 did not have any alterations. Mismatches occurred only in the T3SS needle tip (*ipaD*) to *S. boydii* 183. Among the mismatched nucleotides in *IpaD* gene, three resulted in amino acid substitutions in the C-terminal domain: asparagine 197 by lysine, arginine 224 by aspartic acid, and arginine 239 by lysine (Figure 9 A). Genes *ipaD* and *ipaB* together form a plug in the needle to insert bacterial proteins into the host cell membrane. The *ipaD* C-terminal domain has an essential role in *ipaB* binding to create the translocation pore for phagosome escape and macrophage apoptosis. The 3-D structure of the *ipaD* amino acid-substituted sequence showed a slight modification from the *ipaD* present in *S. flexneri* str. 201 (Figure 9 B). Such *IpaD* gene modifications of *S. boydii* str. 183 could affect its natural binding to *ipaB*, which could be directly associated with invasiveness loss for this bacterium.

**Figure 9.**
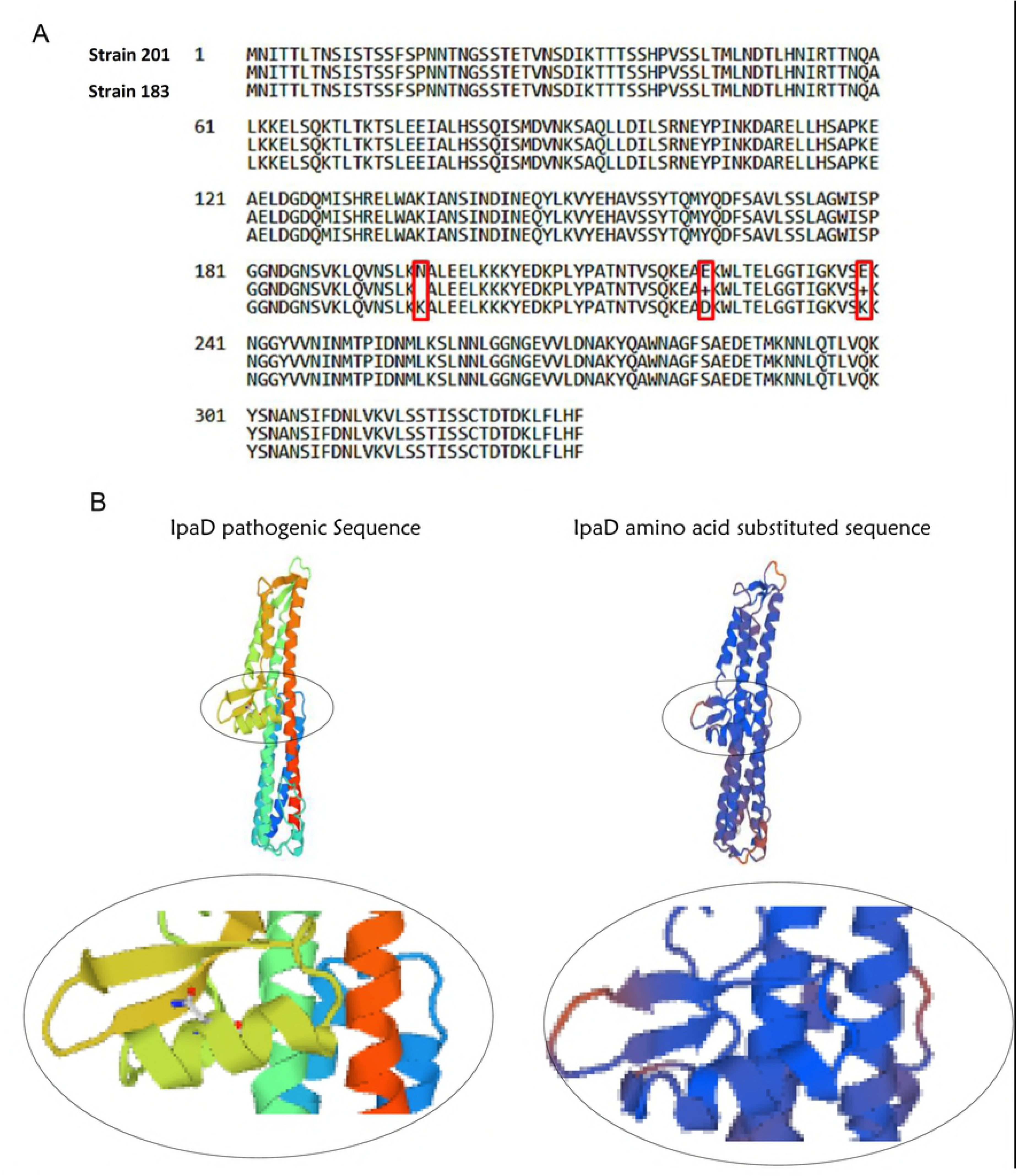
Mismatch occurrence in tip of T3SS apparatus needle. The IpaA, IpaB, IpaC and IpaD genes and amino acid sequences compared for mismatches. A) Only *ipaD* from *Shigella boydii* strain 183 showed three mismatches that resulted in amino acid substitutions in the C-terminal domain: IpaD asparagine 197 by lysine; arginine 224 by aspartic acid; and arginine 239 by lysine. B) Remodeling of these structures *in silico* showed the predicted 3D-structure of *Shigella boydii* strain 183 shows changes in the essential area of the T3SS apparatus needle where IpaB binds IpaD.

## DISCUSSION

Here, we characterized the genetic content of a wild-type *Shigella boydii* and compared its virulence potential at *in vitro* and *in vivo* experiments against wild-type *Shigella flexneri*. Both isolates were from Amazon region diarrheic children (5) and had T3SS-related genes identified by PCR assay (8,9). The presence of *S. boydii* outside of Indian subcontinent is described as 1-2% frequency in the literature (1,3) and the high number of this *Shigella* species found in Amazon was unusual. Clinical *Shigella* genomic studies are needed to elucidate its virulence mechanism and contribute to *Shigella* vaccine development (2). Among *Shigella* species, *S. boydii* is neglected due to its supposed lack of clinical relevance, one of the reasons why there is little known about *S. boydii* virulence determinants and its association with shigellosis (3).

Classical studies show that *Shigella* virulence rely on their ability to invading and colonize the intestinal epithelium, and multiplication of pathogens within the large intestine mucosa is a requirement for bacillary dysentery (21–24). Here, the HEp-2 cell invasion assay was essential to establish the non-invasive phenotype of *S. boydii* str. 183. As expected, *S. boydii* str. 183 was non-invasive compared to virulent *S. flexneri* str. 201 at *in vivo* and *in vitro* assays. Several live attenuated oral *Shigella* vaccines derived from wild-type strains contains precise mutations in the genes encoding enterotoxins (25–27). Interestingly, wild-type *S. boydii* str. 183 appears to be a naturally mutant occurring lacking *set-1A, set-1B*, and *sen/ospD3*. *S. boydii* str. 183 was first isolated from co-infected rotavirus children presenting acute diarrhea but not dysentery. The absence of invasion and enterotoxins by this strain suggest that it has natural attenuating virulence determinants, and *S. boydii* str. 183 was present but was not the pathogen responsible for the diarrheal illness.

Furthermore, while the mechanism by which *Shigella* spp. Invades and spreads within the epithelium is well described, the devices by which bacteria manipulate host cell signaling and the innate immune response are still being determined (28–30). Once adult mice do not exhibit disease symptoms upon oral administration of virulent *Shigella* spp. we chose the pulmonary infection route. Although this model lacks clinical relevance in context to the site of infection, in practice it’s a useful tool to measure immunization efficacy, protection against disease and determine the severity of infection (31). The lung model constitutes an organized mucosal lymphoid organ with T and B lymphocytes and antigen-presenting cells, when *Shigella* is inoculated intranasally occurs acute bronchopneumonia simulating shigellosis (32–38).

We assessed whether our *S*. *boydii* str. 183 was able to subvert the host innate response. Of twenty-one tested innate and adaptive immune response genes, *S. boydii* str. 183 was not able to up-regulate innate immune response genes, in opposition of the virulent *S. flexneri* 201 which presented an inflammatory response (30,32,39,40). Instead, *S. boydii* str. 183 up-regulated TH1-like gene, a novel negative transcription elongation factor involved in transcriptional pausing during cellular migration dynamics, which has a homolog in *Drosophila* TH1. This gene strongly inhibits carboxyl-terminal kinase (PAK1), which is included in cell proliferation and motility (22,41,42). According to previous studies, some Type III Secretion System effector proteins induce intracellular actin remodeling by binding the domain of PAK1 as a virulence strategy to subvert the eukaryotic cell cytoskeleton (43,44). Hence, the ability to induce cell actin remodeling for spreading to adjacent cells is naturally attenuated in this bacteria. Further work is required to elucidate whether up-regulation of TH1-like in *S. boydii* str. 183-infected mice would respond to intracellular actin remodeling and thereafter be unlikely to demonstrate host subversion in *Shigella* infection, as has been established (28,32,45–47).

Still, IFNγ is known to be involved in protection against *Shigella* infections and response to subunit polypeptide vaccines (48). Surprisingly, *S. boydii* str. 183 induced lower IFNγ-positive regulation, and the network revealed that almost all genes were positively correlated with IFN-γ expression (Figure 4). These findings are consistent with the notion that IFN-γ might play a role in both pathogenesis and protection in *Shigella* infection. Meanwhile, the TNF-α driven network of *S. flexneri* str. 201 was coherent with a substantial increase in mRNA abundance for inflammatory mediators and the lethal characteristic of this isolate (10,49–51). Consistent with notion the *S*. *boydii* strain 183 was unable to subvert the host innate response, the virulome-wide analysis showed this bacterium is devoid of the SPATE genes (Pic, SepA, and SigA), described in *Shigella flexneri* as central role in trigger inflammatory response and evolve progression of the disease (Dautin, 2010; Gutiérrez-Jiménez et al., 2008; Ruiz-Perez et al., 2011). Furthermore, it is crucial to emphasize *S. boydii* str. 183 carried only Pic gene. Based on the role of SPATE participation in strong inflammatory response and disease progression, our findings reinforce *S. boydii* str. 183 cannot disturb its host, and could perhaps establish a commensal relationship.

The *Shigella* spp. virulence is dependent on Type III Secretion Systems (T3SS), composed of a needle structure upon an inner rod component assembled on the basal body. The virulence regulators consist of IpaABCD invasion plasmid antigens and mxi-spa-encoded secretion and molecule chaperones, which are indispensable for virulence (1,30,32,52,53). Once the needle is shaped, they form an injection device that delivers effector proteins to host cells following the sensing of host cell contact (54–56). Here, the virulome displayed a complete T3SS apparatus in both bacteria. Moreover, *ipaA*, *ipaB*, *ipaC*, and *ipaD* genes were equally expressed during challenge despite differences in their virulence (Figure 8).

Thus, two findings could elucidate the inability of *S. boydii* str. 183 to multiply into the cytoplasm. First and more plausible, this bacterium did not harbor the pivotal effector *OspE1* and OspE2 to promote bacterial epithelium colonization in both *in vitro* and *in vivo* infectious systems, a requirement for bile-induced bacterial adherence, since a double mutant lost the phenotype (57,58). Second, non-synonymous mismatches occurred only in the *ipaD* C-terminal gene of strain 183, which serves as a scaffold for *ipaB* and many effector proteins. Here, decreased invasion of *S. boydii* str. 183 into the cytoplasm could be assigned to these non-synonymous mismatches, whereas modifications in *ipaD* C-terminal fail to bind the IpaB molecule that is essential for invasion and phagosome escape, as demonstrated with ipaD point mutants (58–60). In turn, exacerbation of *S. flexneri* str. 201 invasiveness has been assigned to some genes harbored by this bacterium, in addition to OspE1 and OspE2 effectors, such as serine-protease autotransporter genes *pic*, *vat*, and *sepA*, as well as *shET1* and *shET2* enterotoxins, suggesting these virulence factors were preponderant for severe lung injury in mouse (1,9,30,32,52,53,61–68).

*Shigella* spp. main virulence determinant it is through Type III Secretion Systems (T3SS) (Killackey et al., 2016). Additional Secretion Systems are found only in a few *Shigella* species (69). In regards to minor Secretion Systems, it’s known the presence of the Type 2 Secretion System (T2SS) in *Shigella dysenteriae* and *Shigella boydii* species (69). The T2SS is encoded by genes of the general secretion pathway (*gsp*) and excretes out the proteins secreted into periplasmic space and is conserved in Gram-negative bacteria (1,70). The *gsp* products from *S. dysenteriae* (Sd197) and *S. boydii* (Sb227) show similarity to the *gsp* genes from enterotoxigenic *Escherichia coli* (ETEC) and *Vibrio cholerae* responsible for secreting the *E.coli* heat labile toxin (Ltx) and cholera toxin (Ctx), respectively. However, at *S. boydii* (sb227), T2SS seems to be inactive due to a frameshift in *gspC* and a nonsense mutation in *gspD* (69).

The presence of Type VI Secretion System (T6SS) in the *Shigella* genus is restricted to *Shigella sonnei* which presents T6SS orthologs (71). T6SS is a syringe-like contractile injection system which perforates prokaryotic or eukaryotic cells and releases toxic effectors proteins directly into the target cells in a single cell-contact dependent step (72) and plays a vital role in competition and virulence (73). Recently, T6SS expression by *S. sonnei*, absent in *S. flexneri*, was associated with a competitive advantage at the host gut in the presence of mixed cultures in a T6SS-dependent manner (73). In commensal bacteria, T6SS play an essential role in defense against invading pathogens and might be one dominant player dictating microbial composition in the host gut. The presence of T6SS in gram-negative pathogens may be an evolutionarily conserved mechanism used by Gram-negative enteric pathogens to establish themselves in the densely populated gut to further cause disease (74). Here our wild-type *Shigella boydii* str. 183 naturally lacking enterotoxins was isolated from a co-infected rotavirus child with acute diarrhea, strongly suggesting that this non-virulent strain was present, but was not the responsible illness pathogen.

Further studies with this strain are necessary to determinate if the presence of T6SS play this role. The competition advantage conferred to *S. sonnei* that the by T6SS against other invasive pathogens (74) associated with the lack of virulence of *S. boydii* str. 183 could be used in probiotic-like future studies.

In conclusion, among *Shigella* species, *S. boydii* is neglected due to its supposed lack of clinical relevance, one of the reasons why there is little known about *S. boydii* virulence determinants and its association with shigellosis (3). Here we profoundly studied the genetic content of *S. boydii* str. 183 and the absence of enterotoxins and the fact of this isolate are initially from co-infected rotavirus children with acute diarrhea, strongly suggests that it was not the responsible illness pathogen. The presence of *S. boydii* outside of Indian subcontinent is described as 1-2% (1,3) and here, we characterized the genetic content of a wild-type *Shigella boydii* isolated from Amazon region (5). The *Shigella boydii* str. 183 virulome was devoid of *shET1* and *shET2* enterotoxins genes, OspE1, OspE2 effectors, and new SPATEs autotransporter genes.

Furthermore, a non-synonymous mutation in the *ipaD* gene could affect the interaction with effector proteins, like the intracellular actin remodeling. This results would explain its inability to multiply in the cytoplasm and subvert the immune system. Still, *S. boydii* str. 183 up-regulated several immune genes demonstrating the ability to elicit an immune response. The considerable shigellosis burden epidemiology motivated us to pursue natural non-virulent *Shigella boydii* that can activate the mucosal immune system without provoking damage in the mucosa. The presence of T6SS could provide a competitive advantage, allowing this non-virulent strain to establish itself at the host gut, promoting an immune response but not the illness. Thus, its non-virulent behavior in our *in vivo* trial could be ascribed as promising at probiotic-like studies.

## Acknowledgments

The author’s thanks to the laboratory technicians of FIOCRUZ/AM, FIOCRUUZ/RJ, FIOCRUZ/RO, INPA, USP/SP, and USP/RP. We thanks the Ph.D. student Allyson Guimarães da Costa to the help in Cytoscape Analysis. And we especially thanks to the DNA Sequencing Platform of FIOCRUZ/IOC through the person of Ana Carolina Vicente de Paulo by the possibility of execution of the DNA sequencing method.

